# Evolution of rarity and phylogeny determine above- and belowground biomass in plant-plant interactions

**DOI:** 10.1101/2023.11.10.566621

**Authors:** Alivia G. Nytko, John K. Senior, Julianne O’Reilly-Wapstra, Jennifer A. Schweitzer, Joseph K. Bailey

**Author notes:** Corresponding Author (AGN).

## Abstract

Rare species are often considered inferior competitors due to occupancy of small ranges, specific habitats, and small local populations. However, the phylogenetic relatedness and rarity level of interacting species in plant-plant interactions are not often considered when predicting the competitive response of rare plants. We used a common garden of 25 species of Tasmanian *Eucalyptus*, varying in rarity to allow us to differentiate the competitive abilities of rare versus common species when grown in mixtures varying in phylogenetic relatedness and rarity. We demonstrate increased biomass production of rare plant species when interacting with genetically intermediate neighbors through synergistic non-additive effects not seen in common species. Additionally, we also find that all plants, regardless of rarity status, maintain 47% greater aboveground and 69% greater belowground biomass when interacting with common species compared to the rarest species. However, species-specific interactions with one particular common species, *E. globulus*, yielded a 97% increase in biomass compared to average biomass yields in other interactions, suggesting the importance of *E. globulus* integration into rare species restoration plantings. These results are important because they suggest that the evolutionary processes driving species rarity and the phylogenetic divergence of traits interact to drive ecological dynamics of plant-plant interactions in non-additive ways. Through the ecological and evolutionary consideration of performance traits, rarity, and species-specific effects, we can more accurately predict plant-plant interaction dynamics varying in rarity and relatedness across the landscape.

## Introduction

Abiotic and biotic factors work together to shape patterns of community co-existence, above- and belowground mutualisms, and range dynamics across global landscapes [1,2]. Although abiotic factors are commonly considered the main drivers of plant productivity, biotic factors, such as plant-plant interactions, also serve as a mechanism of biodiversity maintenance and ecosystem function [3,4]. Through facilitative and competitive interactions, plant-plant interactions serve as a mechanism that affects biodiversity, including plant community composition, functional diversity, resource use, and response to environmental change factors at local and global scales [5]. Plant-plant competition and facilitation are also important interactions driving local adaptation at the edges of species’ ranges through the limitation or enhancement of range expansion and range boundary shape attributable to shifts in growth and dispersal [6–9]. While plant-plant competition and facilitation can impact many important ecological and evolutionary dynamics, we are only beginning to understand how species interactions may vary depending upon intra- and interspecific variation or other evolutionary processes [10–14]. For example, Williams et al. [15] demonstrates that the phylogenetic relatedness of common plant species in mixture can interact with abiotic factors to affect competitive dynamics along a stress gradient in plant-plant interactions. While these results amend the Stress Gradient Hypothesis (SGH) [16] to suggest that evolutionary processes can drive the competitive response of common plant species in increasingly stressful environments, the rarity of interacting species is not considered. Furthermore, the rarity of a species is often judged based on geographic occurrence with little to no consideration of biotic factors in species persistence; such that a rare species is one with a constrained range, specific habitat, and small local populations [17]. Consequently, the causes of increasing rarity are a multifaceted combination of ecological, genetic, and evolutionary factors driving shifts in geographic patterns of occurrence, such as the drought-induced evolution of water-use traits leading to habitat specialization, climate mediated contractions in range size, and shifts in pollinator dynamics leading to decreased genetic diversity in local populations [18–20]. As rarity becomes progressively common, it is critical to examine the competitive response of plant species in community interactions varying in relatedness and rarity to accurately predict future patterns of competition, facilitation, and associated ecosystem function.

Species rarity is an increasingly common phenomenon across the globe and can change species interactions within communities across a wide variety of taxonomic groups [21–23]. While evolutionary processes, such as the divergent evolution of biomass in rare versus common species [24], may play a critical role in mediating the productivity of rare plant species in genetically diverse communities, many studies employ a strictly ecological approach in examining the response of rare species in community interactions [25-27; but see 28]. Consequently, understanding variation in species interactions depending upon the rarity status and phylogenetic relatedness of plant species within a community is understudied and largely unknown. However, a few studies have examined evolutionary patterns of negative frequency dependency in pollination and growth that can act as stabilizing forces in plant-plant interactions [29,30]. Through the enhanced growth and pollination of rare species in the presence of common neighbors, stabilizing forces in community interactions may act as evolutionary rescues reducing intrinsic disadvantages of rare species associated with small ranges, habitats, and populations [29,30]. Similarly, the coupling of species in community mixtures due to shared above- and belowground partners can drive community assembly and subsequent persistence and coexistence of rare and common species [31]. While the mechanisms underlying rare and common species co-existence suggest that evolutionary processes drive patterns of competition and facilitation in plant communities, our study provides an empirical examination of how such evolutionary dynamics explicitly affect the strength and direction of plant-plant interactions varying in both relatedness and rarity.

Although the mechanisms underlying rare plant productivity in community interactions are unknown, phylogenetically similar and dissimilar species display varying facilitative and competitive responses in complex plant communities [32]. Due to limiting similarity, closely related species generally tend to compete more strongly, resulting in decreased fitness [33,34]. Limiting dissimilarity on the other hand, can result in competitive exclusion due to broad differences in resource demand and competitive ability among distantly related species [35,36]. Although general trends are found in phylogenetically similar and diverse communities, the effects of phylogenetic relatedness interact with environmental stress and climatic severity to shape performance traits that alter the outcomes and dynamics of plant-plant interactions [15,37]. Due to the perpetual environmental instability associated with the constrained niches of rare species, rare plants may benefit from increasingly common facilitative interactions with phylogenetically similar and intermediate partners to increase fitness in complex communities [38]. While rare plants may experience increases in biomass production in facilitative interactions with phylogenetically intermediate neighbors, species-specific responses to environmental factors and interacting species can also drive unique trends in the outcomes of plant-plant interactions [39,40]. If the productivity of rare plant species is dependent on the rarity level and phylogenetic distance of interacting species, the phylogenetic divergence of traits among rare versus common species may interact with the evolution of rarity to drive the outcomes of plant-plant interactions with further effects on ecological services that are expected to shift in response to changes in community structure. Consequently, understanding the ecological and evolutionary drivers of above- and belowground trait variation in rare species is critical to understanding the drivers of rare species success in complex communities, as well as maintaining functional diversity and ecosystem services [41].

To understand the eco-evolutionary dynamics underlying plant-plant interactions among rare and common species in a phylogenetic framework, we used a full factorial common garden experiment using 25 species of Tasmanian eucalypts of known phylogenetic relatedness and variation in rarity [24]. Previous work [24] showed a phylogenetic basis to performance traits associated with the major determinants of species rarity in these eucalypt species. Traits, such as above- and belowground biomass, demonstrated divergent evolution within clades, but convergent patterns among clades, providing evidence that species traits that allow for rarity have repeatedly evolved across the phylogeny [24]. This is a significant advance as it indicates that rare species have lower above- and belowground biomass than common species and thus may interact uniquely with other plants. We, therefore, hypothesized: 1) Above- and belowground plant biomass is related to the phylogenetic relatedness of interacting species; 2) Above- and belowground plant biomass is related to the rarity level of interacting plant species. Our results show that rare species have enhanced competitive abilities and synergistic non-additive responses in genetically intermediate relationships as well as in interactions with common plant species. The high potential for using designated plant-plant interactions to increase productivity and performance of rare plant species may allow for the maintenance of functionally unique ecosystem services [42].

## Methods

The 25 species of Tasmanian *Eucalyptus* we used are represented by two subgenera (*Symphyomyrtus* and *Eucalyptus*), four phylogenetic sections (*Maidenaria*, *Aromatica*, *Cineraceae*, *Eucalyptus*), and five series (*Globulares*, *Orbiculares*, *Viminales*, *Seminunicolores*, *Foveolatae*) and dominate much of Tasmanian forests **(S1 Table)**. Ranging vastly in height, growth form, and function, rare and common Tasmanian eucalypts vary widely in above- and belowground biomass, indicating that traits associated with rarity have diverged to employ different evolutionary strategies [24]. As such, Tasmanian eucalypts, and their evolutionary history have gained global interest and a complete phylogeny has been resolved to understand evolutionary drivers of ecological patterns [43,44]. Therefore, the Tasmanian eucalypts provide a unique model system to empirically understand how above- and belowground biomass production is driven by the phylogenetic distance and rarity of interacting species due to the well-established phylogeny, diverse levels of rarity, and high occurrence of interactions between common and rare eucalypts [45].

To measure traits associated with rarity, a full factorial common garden experiment consisting of monocultures and mixtures of different species under varying levels of CO_2_ and Nitrogen (N) fertilization was developed using seeds from each species obtained from one to six maternal trees within a single population **(S1 Fig)**. Each *Eucalyptus* species was grown in a two species mixture consisting of monocultures, as well as phylogenetically similar, intermediate, and distant communities. Monocultures represent relationships absent of phylogenetic distance. After approximately five months of growth, seedlings were harvested and the aboveground and belowground biomass was separated, dried, and weighed (g). Above- and belowground biomass was summed to determine total biomass. See details of this experiment from Senior et al. [43]; data from this paper were recategorized through the addition of rarity levels and reanalyzed using continuous phylogenetic distances to address the hypotheses outlined above. Recategorized data is available at https://doi.org/10.6084/m9.figshare.23611818.v5.

All statistical analyses were performed using R Statistical Software (version 4.2.1, R Core Team 2022). Twenty-five species of native Tasmanian *Eucalyptus* were categorized into seven ordinally ranked levels of rarity based on range size, habitat specificity, and population size in accordance with Rabinowitz [17], Williams & Potts [46] & Nytko et al. [24] **(S2 Fig)**. Each form of rarity was represented by at least one species of Tasmanian *Eucalyptus*. Furthermore, interactions varying in phylogenetic relatedness spanned all rarity levels, such that each mixture type consisted of all possible rarity level combinations (S1 Fig). Levels of rarity were assigned to both the target species and interacting species within community mixtures; therefore, they shall be referred to as target rarity level and interacting rarity level respectively. Additionally, variation in growth form among target species was represented by differences in mean adult height (m) and accounted for as a random variable in all analyses. Linear mixed models (LMM) were first used to examine the effects of phylogenetic relatedness, interacting rarity level, CO_2_ addition, and N fertilizer addition on total biomass (“lmer” function in “lme4” package, R). The LMM yielded no interactive effects between phylogenetic relatedness, interacting rarity level, and other fixed effects **(S2 Table)**. Additionally, when accounting for CO_2_ addition and N fertilizer addition as random effects, the relationship between phylogenetic relatedness/interacting rarity level and total biomass remained the same. Therefore, statistical differences in the above- and belowground biomass of eucalypts in different community mixtures varying in phylogenetic relatedness and among different interacting rarity levels were examined using linear mixed models.

To determine the effects of phylogenetic relatedness on the total biomass of target species, the phylogenetic distance between each species was calculated across the phylogeny (“cophenetic.phylo” function in “ape” package, R). Furthermore, the singular and interactive effects of phylogenetic distance and interacting rarity level on total, aboveground, and belowground biomass were analyzed using linear mixed models. Differences in observed mean biomass and expected mean biomass (based on biomass production in monocultures) were calculated for each species as raw [observed-expected] interaction strengths and standardized using target species grouped z-scores. To determine the significance and direction of non-additive effects among species mixtures across all levels of target rarity, standardized interaction strengths were used in one-sample T tests (mu = 0) and linear mixed models. Positive interaction strengths in community mixtures represent synergistic non-additive effects indicative of facilitation, negative interaction strengths represent antagonistic non-additive effects indicative of competition, and neutral interaction strengths represent additive effects indicative of neutral plant-plant interactions [47]. To determine species-specific effects on neighboring biomass, linear mixed models were used to determine the effect of neighboring species on the above- and belowground biomass of target species while accounting for the rarity level and adult mean height of target species as well as the phylogenetic relatedness of interacting species as random variables. Tukey HSD post-hoc analyses were completed for all significant results (“ghlt” and “cld” functions in “multcomp” package, R).

## Results

Overall, the phylogenetic distance and rarity status of interacting neighbor species were strong determinants of above- and belowground biomass (**Table 1**). Eucalypts displayed significantly increased total and belowground biomass production when grown with intermediately related neighboring species (**Fig 1a**; Table 1). Specifically, exceptionally rare species experienced a significant increase in total biomass when interacting with intermediately related species, not seen in common counterparts (**Fig 1b**). For example, species belonging to rarity level two experienced a 155% increase when interacting with intermediately related neighbors compared to monocultures and a 57% increase compared to distantly related neighbors. In contrast, common species had high biomass irrespective of the neighboring plant environment (Fig 1b). Importantly, irrespective of the phylogenetic distance of the interacting species, there were non-additive synergistic effects where rare species over-performed in mixture based upon expectations when grown in monoculture (**Fig 2**; **Table 2**). Moreover, the strength and direction of the performance of rare species depended upon their rarity level (Table 2). For example, the interaction strengths of species rarity level 1 and 2 were strongly positive compared to rarity level 7 which varied from positive to negative and behaved similarly to common species (Fig 2). Common species generally exhibited absent or negative (i.e., antagonistic) non-additive effects. Specifically, common species demonstrated an average standardized interaction strength of 0.02 compared to an interaction strength of 0.463 in the rarest species: a difference of 183%. Variation in non-additive effects of species mixtures between rare and common species was most evident in closely and intermediately related species; demonstrated by an interaction strength of 0.450 in the rarest species versus an interaction strength of 0.080 in common species within closely related mixtures, and an interaction strength of 0.856 in rarity level two species versus an interaction strength of -0.893 in common species within intermediately related mixtures (Fig 2b).

**Table 1.** Results of linear mixed models, examining the singular and interactive effects of interacting rarity level (1-7 and common) and phylogenetic distance on total, aboveground, and belowground biomass. Variation in mean adult height (m) was accounted for as a random variable. Alpha = 0.05.

| Response | Effects | DF | Chisq | P value |
| --- | --- | --- | --- | --- |
| Total Biomass | Interacting Rarity | 7 | 41.944 | 5.33e-07* |
|  | Phylogenetic Distance | 1 | 4.002 | 0.045* |
|  | Interacting Rarity x<br>Phylogenetic Distance | 7 | 12.138 | 0.096 |
| Aboveground<br>Biomass | Interacting Rarity | 7 | 42.696 | 3.819-07* |
|  | Phylogenetic Distance | 1 | 3.759 | 0.053 |
|  | Interacting Rarity x<br>Phylogenetic Distance | 7 | 11.362 | 0.124 |
| Belowground<br>Biomass | Interacting Rarity | 7 | 36.260 | 6.474e-06* |
|  | Phylogenetic Distance | 1 | 3.973 | 0.046* |
|  | Interacting Rarity x<br>Phylogenetic Distance | 7 | 12.959 | 0.073 |

**Fig 1.**
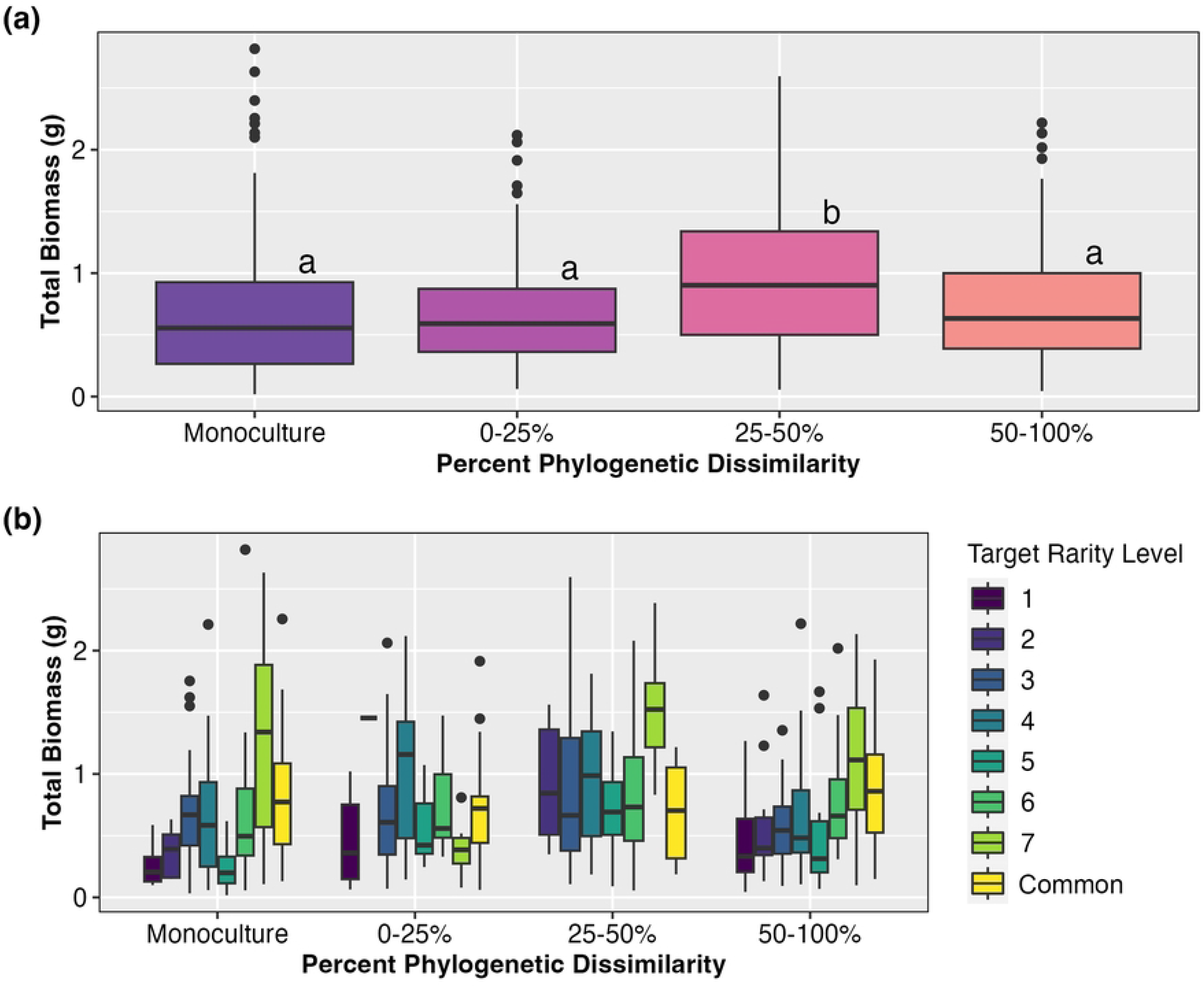
a) Comparative boxplots demonstrating the total biomass of 25 species of Tasmanian *Eucalyptus*, varying in rarity by mixture representative of phylogenetic relatedness and b) variation in standardized interaction strengths representative of synergistic and antagonistic non-additive effects. Monocultures represent an absence of phylogenetic distance. Positive standardized interaction strengths represent synergistic non-additive effects, while negative standardized interaction strengths represent antagonistic non-additive effects.

**Fig 2.**
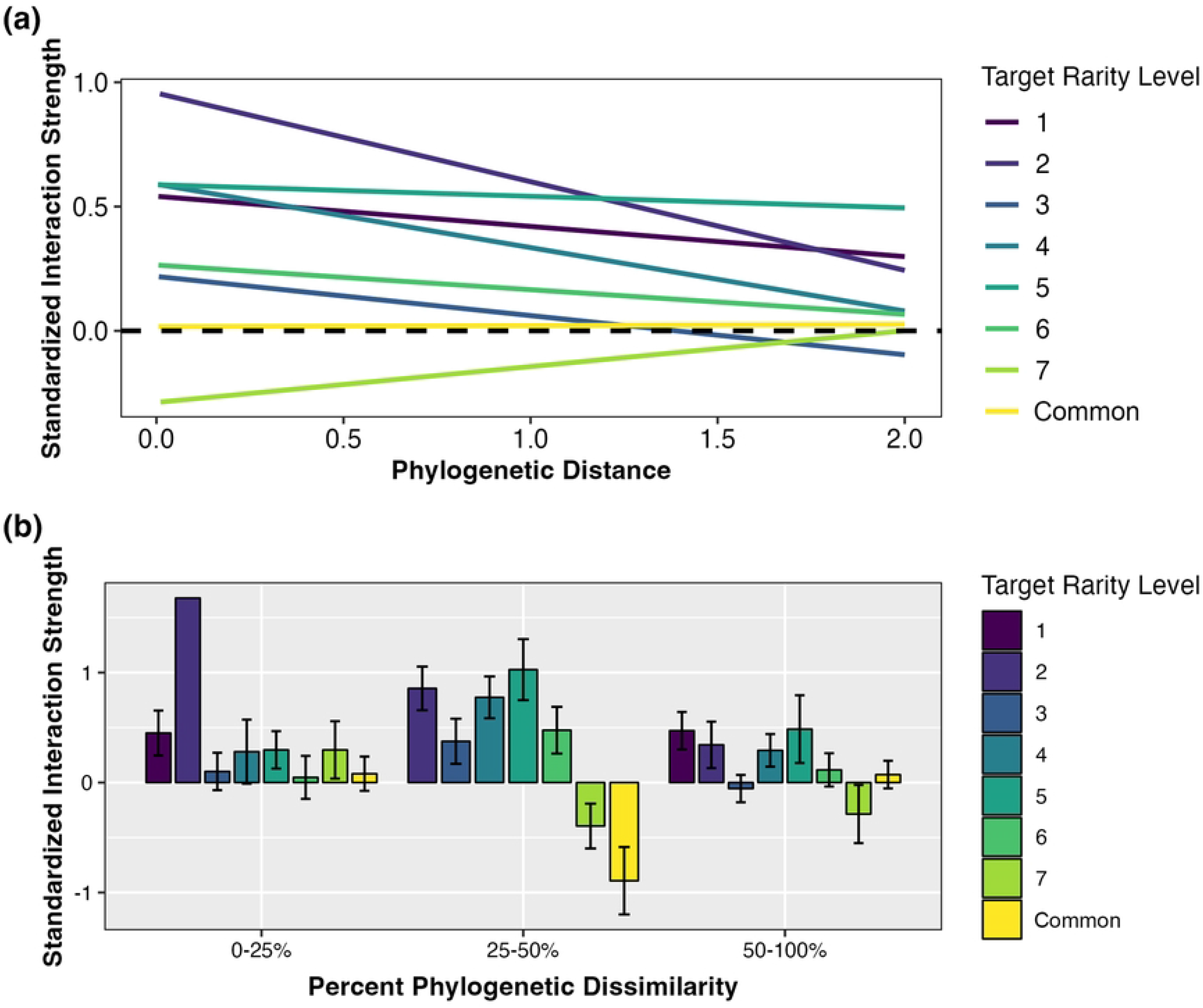
Linear regressions and comparative boxplots demonstrating the negative relationship between phylogenetic distance and standardized interaction strengths by target rarity level.

**Table 2.** a) Results of one-sample t-test (mu = 0) examining the difference between observed biomass in mixtures and the expected biomass based on biomass production in monocultures by target rarity level. A significant divergence of standardized interaction strength from the null demonstrates that biomass in mixtures are significantly different than the expectation in monocultures. **b) Results of linear mixed model examining the effects of interacting rarity level (1-7 and common) and phylogenetic distance on standardized interaction strengths.** Variation in mean adult height (m) was accounted for as a random variable. Alpha = 0.05.

| a) One Sample t-tests |  |  |  |  |
| --- | --- | --- | --- | --- |
| Response | Target rarity level | DF | t-value | P value |
| Standardized Interaction Strengths | 1 | 45 | 3.930 | 2.895e-04* |
|  | 2 | 23 | 4.321 | 2.529e-04* |
|  | 3 | 102 | 1.702 | 0.092 |
|  | 4 | 63 | 3.676 | 4.914e-04* |
|  | 5 | 26 | 3.673 | 0.001* |
|  | 6 | 82 | 2.506 | 0.014* |
|  | 7 | 42 | -2.262 | 0.029* |
|  | Common | 104 | -0.665 | 0.507 |
| b) Analysis of Variance |  |  |  |  |
| Response | Effects | DF | Chisq | P value |
| Standardized Interaction Strengths | Interacting Rarity Level | 7 | 23.787 | 0.001* |
|  | Phylogenetic Distance | 1 | 5.677 | 0.017* |
|  | Interacting Rarity x Phylogenetic Distance | 7 | 11.638 | 0.113 |

Across all phylogenetic backgrounds, the rarity level of interacting species had a significant effect on the total, aboveground, and belowground biomass of all target species. Regardless of target rarity level, species maintained an average of 47% greater aboveground and 69% greater belowground biomass when interacting with the most common species compared to the rarest species (**Fig 3a**; Table 1). Communities containing common neighbors had higher biomass than communities comprised of only rare species, given equal abundance in the community. Further, communities of phylogenetically intermediate distance experienced the greatest impact on total biomass from common neighbors (**Fig 3b**). However, as target rarity and interacting rarity increased, biomass generally increased.

**Fig 3.**
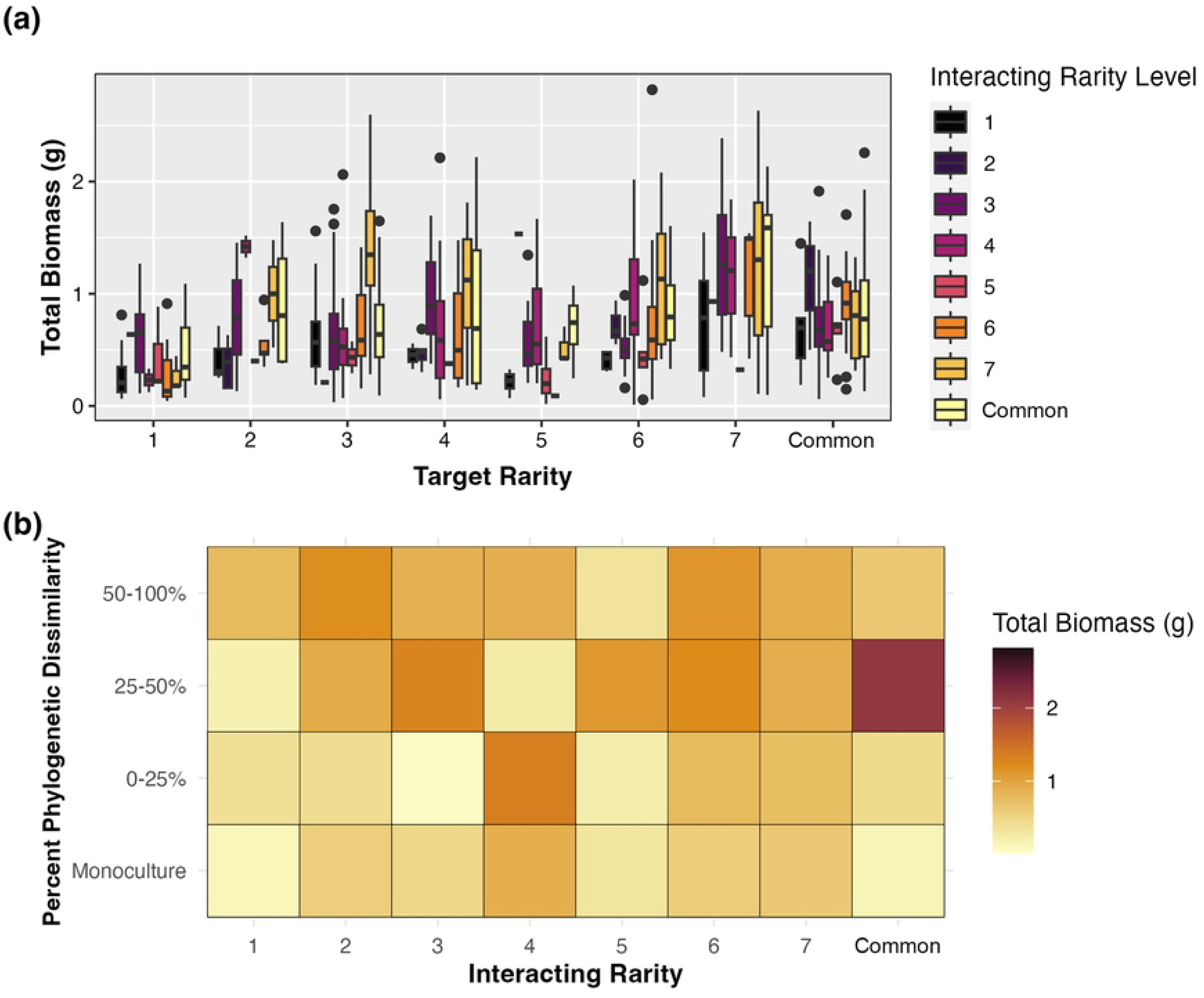
Comparative boxplots and heat map demonstrating the relationship between the interacting rarity level and the phylogenetic distance of neighboring plant species on total biomass.

Finally, we were surprised to find that certain interacting species displayed positive and negative species-specific effects on the biomass of neighboring species regardless of rarity or phylogenetic relatedness (**Fig 4**; **Table 3**). While more common interacting species tended to increase the biomass of rare neighbors, select species (*E. brookeriana*, *E. globulus*, and *E. ovata*) significantly increased the productivity of all neighboring species (**Table 4**). Specifically, when planted with *E. globulus*, target species displayed a 97% increase in biomass compared to interactions with other species irrespective of phylogenetic relatedness. On the other hand, eight species (*E. perriana*, *E. pulchella*, *E. radiata*, *E. regnans*, *E. rubida*, *E. subcrenulata*, *E. tenuiramis*, and *E. vernicosa*) had significant inhibitory effects on the biomass of target neighbor species and reduced growth by 31-54% (Table 4).

**Fig 4.**
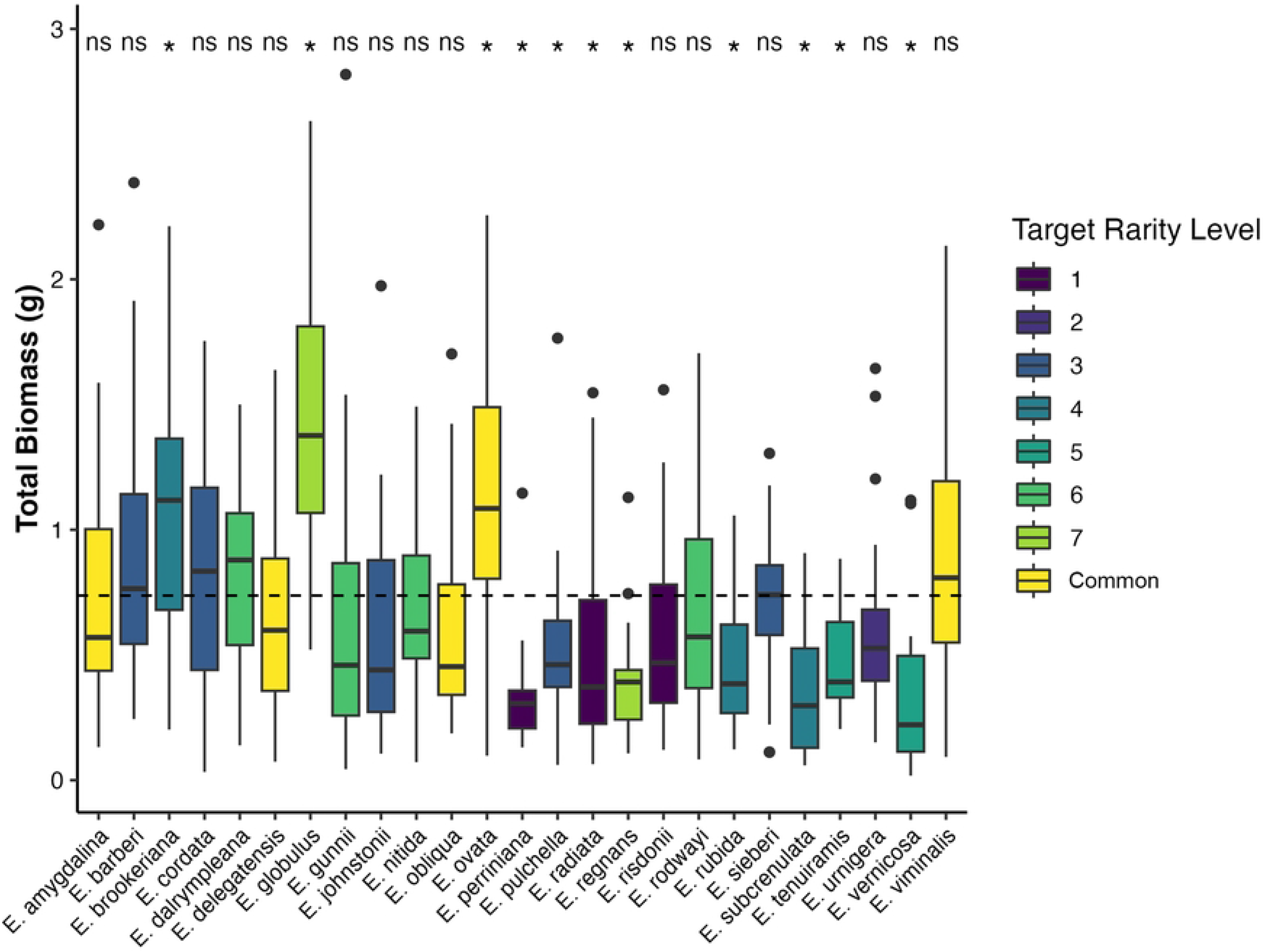
Comparative boxplots demonstrating species-specific effects of interacting species on target species biomass production. The dashed line signifies mean total biomass.

**Table 3.** Results of linear mixed model examining the effect of interacting species on the total, aboveground, and belowground biomass of target species. Target rarity level, adult mean height, and phylogenetic relatedness of interacting species was accounted for in the error structure as random variables. Alpha = 0.05.

| Response | DF | Chisq | P value |
| --- | --- | --- | --- |
| Total biomass | 24 | 221.09 | <2.2e-16* |
| Aboveground biomass | 24 | 226.81 | <2.2e-16* |
| Belowground biomass | 24 | 176.52 | <2.2e-16* |

**Table 4.**
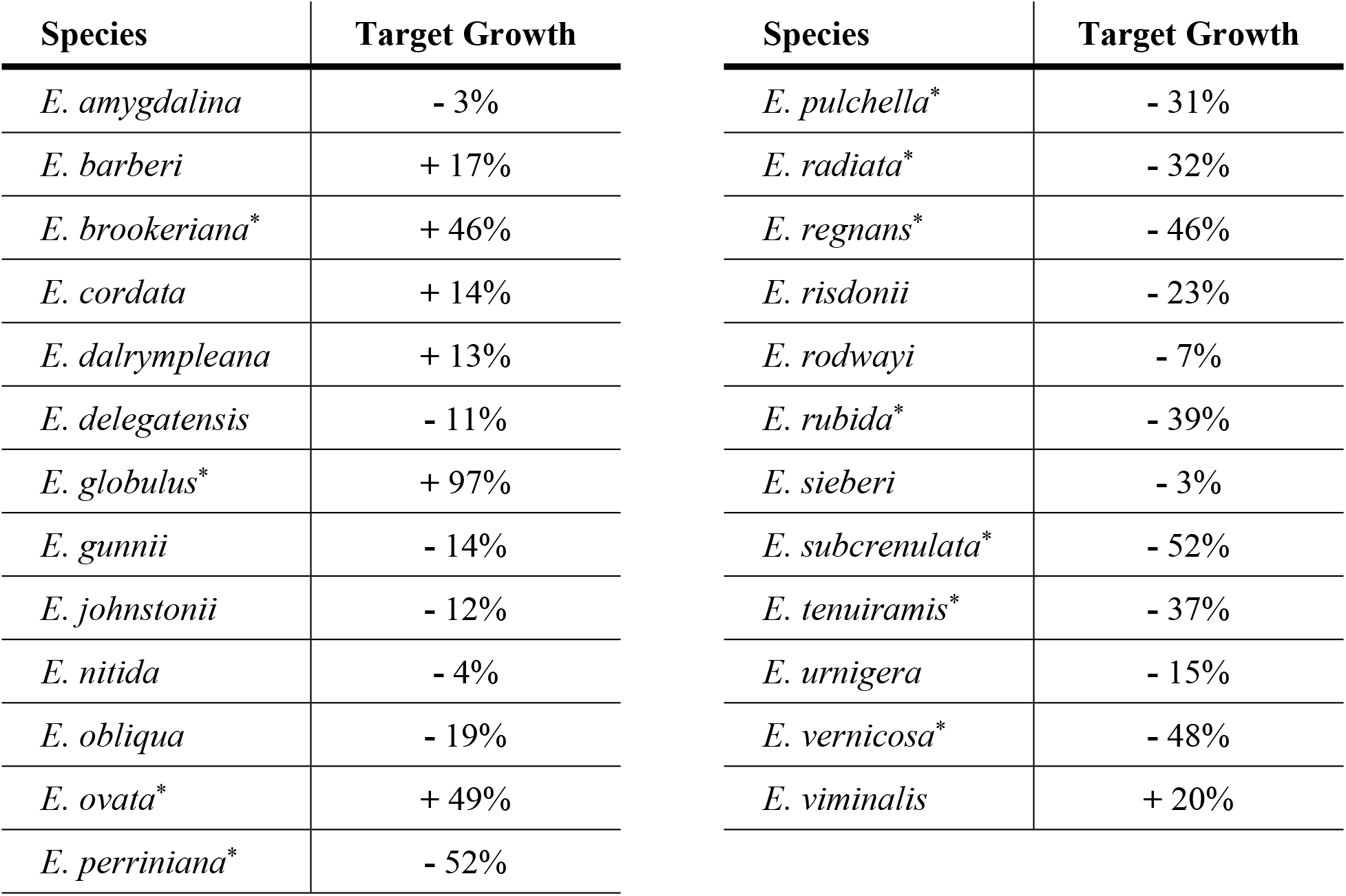
Summary average of species-specific effects of neighboring species on target species biomass across all interaction types. Positive target growth values represent increased biomass of neighboring species, while negative target growth values represent decreased biomass of neighboring species compared to the mean biomass of all target species across all mixture types. Alpha = 0.05.

| Species | Target Growth | Species | Target Growth |
| --- | --- | --- | --- |
| *E. amygdalina* | - 3% | *E. pulchella* * | - 31% |
| *E. barberi* | + 17% | *E. radiata* * | - 32% |
| *E. brookeriana* * | + 46% | *E. regnans* * | - 46% |
| *E. cordata* | + 14% | *E. risdonii* | - 23% |
| *E. dalrympleana* | + 13% | *E. rodwayi* | - 7% |
| *E. delegatensis* | - 11% | *E. rubida* * | - 39% |
| *E. globulus* * | + 97% | *E. sieberi* | - 3% |
| *E. gunnii* | - 14% | *E. subcrenulata* * | - 52% |
| *E. johnstonii* | - 12% | *E. tenuiramis* * | - 37% |
| *E. nitida* | - 4% | *E. urnigera* | - 15% |
| *E. obliqua* | - 19% | *E. vernicosa* * | - 48% |
| *E. ovata* * | + 49% | *E. viminalis* | + 20% |
| *E. perriniana* * | - 52% |  |  |

## Discussion

The rarity of plants in a community is often considered a consequence of random stochastic events that shape the potential range of a species [48]. However, recent work demonstrates that performance traits like biomass production can be strong determinants of range size and habitat specificity that are related to the rarity of species [24,49,50]. Building upon our previous work demonstrating an evolutionary basis to traits conferring plant rarity [24], this study shows that the above- and belowground biomass of plants can vary depending upon the rarity of the interacting species and the phylogenetic relatedness between those species. Our results run counter to the prevalent hypothesis that species rarity is representative of one step closer to extinction and instead suggest that rare species may utilize unique plant-plant interactions to persist in a rare state. These results are important because they support analogous lines of research to demonstrate that evolved traits can determine the outcomes of species interactions in non-additive ways with further impacts on conservation and restoration efforts.

Controlling for phylogenetic relatedness and the evolutionary non-independence of interacting plant species has been shown to be important in biodiversity and ecosystem function relationships [51,52]. For example, Cadotte et al. [53] and Verdú et al. [32] demonstrated the importance of underlying phylogenetic signals and spatial patterns of phylogenetic relationships on the functional traits that mediate community composition and stability across the landscape. However, we still lack a good understanding of how phylogenetic relatedness impacts plant-plant interactions and ultimately plant performance. This is largely due to the difficulty in manipulating phylogenetic relatedness in an experimental framework [but see 12,54]. Consistent with previous research, our study experimentally demonstrates that the phylogenetic relatedness of interacting species also plays a central role in driving the biomass production of rare species in plant-plant interactions. The demonstrated increase in biomass production of rare species when interacting with phylogenetically similar and intermediate species suggests that rare species may escape the negative effects of competition under limiting similarity [55,56]. One hypothesis for these patterns is that associations with phylogenetically similar and intermediate neighbors allows for the expansion of realized niche space through the differentiation of resource use, sharing of mutualistic partners, and evasion of competitors [41,42,57,58]. Plant species often demonstrate coevolutionary patterns between functional traits and mechanisms of resource uptake, resource use, and pollination systems [59–65]. Through facilitative interactions with closely and intermediately related species, rare species may engage with and benefit from the specialized systems of resource acquisition and pollination in genetically similar neighbors that may go unrecognized and unused in distantly related species.

These results also show that the evolution of performance traits that drive patterns of rarity are important factors determining the strength and direction of plant-plant interactions. Historically, rare species have been considered inferior competitors when compared to more common species [50,66,67]. However, our results suggest that the divergence of performance traits in rare versus common species across the phylogeny can drive the response of species in plant-plant interactions independently from phylogenetic relatedness. While common species maintain higher levels of above- and belowground biomass [24,49,50], the effect of plant-plant interactions on the productivity of rare species is more nuanced than originally hypothesized [66]. While competitive interactions are common, facilitative relationships between rare and common species that increase the biomass of the rare partner, also occur, and may increase in frequency with increasing environmental stress [41,42,57,58]. For example, the variation in non-additive responses of rare and common species in mixed environments suggests that the interplay between rare and common species is more complicated than basic competition. One hypothesis for these patterns is that rare-common plant interactions may be mediated through shared microbial communities, altered biochemical and physiological patterns of defense, and shifts in herbivory associated with highly productive common neighbors that allow for niche expansion and increased biomass production that would otherwise be unattainable for isolated rare species in monocultures [27,69–71]. One interesting question that arises from these results is: can rare plant species competitively exclude common species when interacting with closely and intermediately related neighbors due to synergistic non-additive effects on performance? Prevalent non-additive synergistic effects mediating the potential dominance of rare plant species in community dynamics may be attributed to functionally unique traits that allow for the inhabitation of stressful environments through association with beneficial soil microbial communities, enhanced pollination, and use of undesirable niche space [37,72,73]. Furthermore, the small geographic and evolutionary breadth in which rare plant species persist within competitive communities suggests that restricted range, habitat, and populations may be caused by the niche restriction of community interactions. Consequently, rare species interacting with phylogenetically similar and intermediately related species may utilize non-additive responses to expand their niche space and dominance within communities across the landscape [74].

A potential mechanism underlying the increased biomass associated with common species lies in the strong relationships that exist between dominant dual mycorrhizal plantation species and soil microbial communities that sequester, and share acquired resources with interacting neighbors [75]. Although many species involved in the beneficial species-specific interactions demonstrated in this study vary in growth form, nutritional acquisition, microbial symbionts, and conservation status, rare species may benefit from interactions with not only common species but also specific species such as *E. globulus*. Although the mechanisms underlying facilitative interactions involving *E. globulus* are not fully understood, *E. globulus* can enhance nutrient and water availability in community mixtures, creating a favorable environment for neighboring plant species [76,77]. Therefore, rare species conservation can be significantly improved through the incorporation of highly productive species associated with increased biomass of rare species in preexisting restoration plantings. Consequently, increases in rare species persistence and productivity may be both economically and environmentally favorable through the augmentation of functional diversity, rare species biomass, and ecosystem function. Therefore, through the consideration of evolved performance traits, in addition to species-specific effects, patterns of plant-plant interaction dynamics involving rare and common species can be more accurately predicted and accounted for in conservation efforts across the landscape.

## Data Availability

The data that support the findings of this study are openly available in figshare at https://doi.org/10.6084/m9.figshare.23611818.v5. The data were also used to support the findings of Senior et al. [29] available at https://doi.org/10.1371/journal.pone.0060088.

## Acknowledgements

We thank the National Science Foundation for funding through the Graduate Research Fellowship (GRFP) to AGN (award number: 2022341820). We are also grateful to Hannah Shulman for assistance with figures and data visualization.

## Author Contributions

Conceived and designed the experiment: JKB JAS JO. Performed the experiment: JKS. Analyzed the data: AGN. Wrote the manuscript: AGN. Edited and revised the manuscript: AGN JKB JAS JKS JO.

## References Cited

1. Jeffers ES, Bonsall MB, Froyd CA, Brooks SJ, Willis KJ. The relative importance of biotic and abiotic processes for structuring plant communities through time. J. Ecol. 2015; 103(2): 459–472.

2. Wisz MS, Pottier J, Kissling WD, Pellissier L, Lenoir J, Damgaard C, et al. The role of biotic interactions in shaping distributions and realised assemblages of species: implications for species distribution modeling. Biol. Rev. Camb. Philos. Soc. 2013: 88(1): 15–30.

3. Belovsky GE. & Slade JB. Biotic versus abiotic control of primary production identified in a common garden experiment. Sci. Rep. 2019; 9, 11961.

4. Bashirzadeh M, Soliveres S, Farzam M, Ejtehadi H. Plant-plant interactions determine taxonomic, functional and phylogenetic diversity in severe ecosystems. Glob. Ecol. Biogeogr. 2022; 31(4): 649–662.

5. Brooker RW. Plant-plant interactions and environmental change. New Phytol. 2006; 171(2): 271–284.

6. Brooker RW, Maestre FT, Callaway RM, Lortie CL, Cavieres LA, Kunstler G, et al. Facilitation in plant communities: the past, the present, and the future. J. Ecol. 2008; 96(1): 18–34.

7. Legault G, Bitters ME, Hastings A, Melbourne BA. Interspecific competition slows range expansion and shapes range boundaries. Proc. Natl. Acad. Sci. U.S.A. 2020: 117(43): 26854–26860.

8. Michalet R, Le Bagousse-Pinguet Y, Maalouf JP, Lortie CJ. Two alternatives to the stress gradient hypothesis at the edge of life: the collapse of facilitation and switch from facilitation to competition. J. Veg. Sci. 2013; 25(2): 609–613.

9. Lyu S. & Alexander JM. Competition contributes to both warm and cool range edges. Nat. Commun. 2022; 13(1), 2502.

10. Genung MA, Bailey JK, Schweitzer JA. Welcome to the neighbourhood: interspecific genotype by genotype interactions in *Solidago* influence above and belowground biomass and associated communities. Ecol. Lett. 2012; 15(1): 65–73.

11. Genung MA, Bailey JK, Schweitzer JA. Belowground interactions shift the relative importance of direct and indirect genetic effects. Ecol. Evol. 2013; 3(6): 1692–1701.

12. Genung MA, Schweitzer JA, Bailey JK. Evolutionary history determines how plant productivity responds to phylogenetic diversity and species richness. PeerJ. 2014; 2, e288.

13. Kinlock NL. A meta-analysis of plant interaction networks reveals competitive hierarchies as well as facilitation and intransitivity. Am. Nat. 2019; 194(5): 640–653.

14. Soliveres S, Maestre FT, Berdugo M, Allan E. A missing link between facilitation and plant species coexistence: nurses benefit generally rare species more than common ones. J Ecol. 2015; 103(5): 1183–1189.

15. Williams EW, Zeldin J, Semski WR, Hipp AL, Larkin DJ. Phylogenetic distance and resource availability mediate direction and strength of plant interactions in a competition experiment. Oecologia. 2021; 197: 459–469.

16. Bertness MD. & Callaway R. Positive interactions in communities. Trends Ecol. Evol. 1994; 9(5): 191–193.

17. Rabinowitz D. Seven forms of rarity, In: Synge, H. (Ed.). The Biological Aspects of Rare Plants Conservation. Wiley, New York; 1981; pp. 205-217.

18. Cavender-Bares J, Kitajima K, Bazzaz FA. Multiple trait associations in relation to habitat differentiation among 17 Floridian Oak species. Ecol. Monogr. 2004; 74(4): 635–662.

19. Yesuf GU, Brown KA, Walford NS, Rakotoarisoa SE, Rufino MC. Predicting range shifts for critically endangered plants: Is habitat connectivity irrelevant or necessary? Biol. Conserv. 2021; 256, 109033.

20. Feigs JT, Holzhauer SIJ, Huang S, Brunet J, Diekmann M, Hedwall P, et al. Pollinator movement activity influences genetic diversity and differentiation of spatially isolated populations of clonal forest herbs. Front. Ecol. Evol. 2022; 10.

21. Enquist BJ, Feng X, Boyle B, Maitner B, Newman EA, Jørgensen PM, et al. The commonness of rarity: Global and future distribution of rarity across land plants. Sci. Adv. 2019; 27, 5(11).

22. Hooper DU, Adair EC, Cardinale BJ, Byrnes JEK, Hungate BA, Matulich KL, et al. A global synthesis reveals biodiversity loss as a major driver of ecosystem change. Nature. 2012; 486: 105–108.

23. Calatayud J, Andivia E, Escudero A, Melián CJ, Bernado-Madrid R, Stoffel M, et al. Positive associations among rare species and their persistence in ecological assemblages. *Nat*. Ecol. Evol. 2020; 4: 40–45.

24. Nytko AG, Senior JK, Wooliver RC, O’Reilly-Wapstra J, Schweitzer JA, Bailey JK. An evolutionary case for rarity. [Preprint]. 2023. Available from: 10.21203/rs.3.rs-3369472/v1. In review at Ecol. Evol.

25. Gibson RH, Nelson IL, Hopkins GW, Hamlett BJ, Memmott J. Pollinator webs, plant communities and the conservation of rare plants: arable weeds as a case study. J. Appl. Ecol. 2006; 43(2): 246–257.

26. Levine JM, McEachern AK, Cowan C. Do competitors modulate rare plant response to precipitation change? Ecol. 2010; 91(1): 130–140.

27. Sritharan MS, Scheele BC, Blanchard W, Lindenmayer DB. Spatial associations between plants and vegetation community characteristics provide insights into the processes influencing plant rarity. PloS One, 2021; 16(12), e0260215.

28. Kempel A, Rindisbacher A, Fischer M, Allan E. Plant soil feedback strength in related to large-scale plant rarity and phylogenetic relatedness. Ecol. 2018; 99(3): 597–606.

29. Bergamo PJ, Streher NS, Traveset A, Wolowski M, Sazima M. Pollination outcomes reveal negative density-dependence coupled with interspecific facilitation among plants. Ecol. Lett. 2019; 23(1): 129–139.

30. Yamamichi M, Kyogoku D, Iritani R, Kobayashi K, Takahashi Y, Tsurui-Sato K, et al. Intraspecific adaptation load: A mechanism for species coexistence. Trends Ecol. Evol. 2020; 35(10): 897–907.

31. Albrecht J, Bohle V, Berens DG, Jaroszewicz B, Selva N, Farwig N. Variation in neighbourhood context shapes frugivore mediated facillitation and competition among co-dispersed plant species. J. Ecol. 2015l 103(2): 526–536.

32. Verdú M, Jordano P, Valiente-Banuet A. The phylogenetic structure of plant facilitation networks changes with competition. J. Ecol. 2010; 98(6): 1454–1461.

33. Violle C, Nemergut DR, Pu Z, Jiang L. Phylogenetic limiting similarity and competitive exclusion. Ecol. Lett. 2011; 14(8): 782–787.

34. Yang Z, Powell JR, Zhang C, Du G. The effect of environmental and phylogenetic drivers on community assembly in an alpine meadow community. Ecology. 2012; 93(11): 2321–2328.

35. Ågren GI. & Fagerström T. Limiting dissimilarity in plants: Randomness prevents exclusion of species with similar competitive abilities. Oikos. 1984; 43(3): 369–375.

36. Kalmykov LV. & Kalmykov VL. A solution to the dilemma ‘limiting similarity vs. limiting dissimilarity’ by a method of transparent artificial intelligence. Chaos Solit. 2021; 146, 110814.

37. Pistón N, Armas C, Schöb C, Macek P, Pugnaire FI. Phylogenetic distance among beneficiary species in a cushion plant species explains interaction outcome. Oikos. 2015; 124(10): 1354–1359.

38. Sánchez-Martín R, Verdú M, Montesinos-Navarro A. Phylogenetic and functional constraints of plant facilitation rewiring. Ecol. 2023; 104(2), e3961.

39. Soliveres S, Eldridge DJ, Hemmings F, Maestre FT. Nurse plant effects on plant species richness in drylands: the role of grazing, rainfall and species specificity. Perspect. Plant Ecol. Evol. Syst. 2012; 14(6): 402–410.

40. Soliveres S. & Maestre FT. Plant-plant interactions, environmental gradients and plant diversity: a global synthesis of community-level studies. Perspect. Plant Ecol. Evol. Syst. 2014; 16(4): 154–163.

41. Jain M, Flynn DFB, Prager CM, Hart GM, DeVan CM, Ahrestani FS, et al. The importance of rare species: a trait-based assessment of rare species contributions to functional diversity and possible ecosystem function in tall-grass prairies. Ecol. Evol. 2014; 4(1): 104–112.

42. Dee LE, Cowles J, Isbell F, Pau S, Gaines SD, Reich PB, et al. When do ecosystem services depend on rare species? Trends Ecol. Evol. 2019; 34(8): 746–758.

43. Senior JK, Schweitzer JA, O’Reilly-Wapstra J, Chapman SK, Steane D, Langley A, et al. Phylogenetic Responses of Forest Trees to Global Change. PLoS One. 2013; 8(4), e60088.

44. Wooliver RC, Marion ZH, Peterson CR, Potts BM, Senior JK, Bailey JK, et al. Phylogeny is a powerful tool for predicting plant biomass responses to nitrogen enrichment. Ecology. 2017; 98(8): 2120–2132.

45. Pfeilsticker TR, Jones RC, Steane DA, Harrison PA, Vaillancourt RE, Potts BM. Expansion of the rare *Eucalyptus risdonii* under climate change through hybridization with a closely related species despite hybrid inferiority. Ann. Bot. 2022; 129: 1–14.

46. Williams KJ. & Potts BM. The natural distribution of *Eucalyptus* species in Tasmania. Tasforests. 1996; 8: 39–149.

47. Michel P, Lee WG, During HJ, Cornelissen JHC. Species traits and their non-additive interactions control the water economy of bryophyte cushions. J. Ecol. 2012; 100(1): 222–231.

48. Zhang X, Pu Z, Li Y, Han XG. Stochastic processes play more important roles in driving the dynamics of rarer species. *J*. Plant Ecol. 2015; 9(3), rtv058.

49. Kempel A, Vincent H, Prati D, Fischer M. Context dependency of biotic interactions and its relation to plant rarity. Divers. Distrib. 2020; 26(6): 758–68.

50. Vincent H, Bornand CN, Kempel A, Fischer M. Rare species perform worse than widespread species under changed climate. Biol. Conserv. 2020; 246.

51. Flynn DFB, Mirotchnick N, Jain M, Palmer MI, Naeem S. Functional and phylogenetic diversity as predictors of biodiversity-ecosystem-function relationships. Ecology. 2011; 92(8): 1573–1581.

52. Joly S, Flynn DFB, Wolkovich EM. On the importance of accounting for intraspecific genomic relatedness in multi-species studies. Methods Ecol. Evol. 2019; 10(7): 994–1001.

53. Cadotte MW, Dinnage R, Tilman D. Phylogenetic diversity promotes ecosystem stability. Ecology. 2012; 93(sp8): S223–S233.

54. Cadotte MW, Albert CH, Walker SC. The ecology of differences: assessing community assembly with trait and evolutionary distances. Ecol. Lett. 2013; 16(10): 1234–1244.

55. Scheffer M. & van Nes EH. Self-organized similarity, the evolutionary emergence of groups of similar species. Proc. Natl. Acad. Sci. U.S.A. 2006; 103(16): 6230–6235.

56. Yan B, Zhang J, Liu Y, Li Z, Huang X, Wanqin Y, et al. Trait assembly of woody plants in communities across sub-alpine gradients: Identifying the role of limiting similarity. J Veg. Sci. 2012; 23(4): 698–708.

57. Basile M. Rare species disproportionally contribute to functional diversity in managed forests. Sci. Rep. 2022; 12, 5897.

58. Leitão RP, Zuanon J, Villéger S, Williams SE, Baraloto C, Fortunel C, et al. Rare species contribute disproportionately to the functional structure of species assemblages. Proc. Royal Soc. B. 2016; 283(1828), 20160084.

59. Heath KD. The coevolutionary genetics of plant-microbe interactions. New Phytol. 2008; 180(2): 268–270.

60. Johnson SD. & Anderson B. Coevolution between food-rewarding flowers and their pollinators. Evol.: Educ. Outreach. 2010; 3: 32–39.

61. Lomáscolo SB, Giannini N, Chacoff NP, Castro-Urgal R, Vázquez DP. Inferring coevolution in a plant-pollinator network. Oikos. 2018; 128(6): 775–789.

62. Lyu D, Msimbira LA, Nazari M, Antar M, Pagé A, Shah A, et al. The coevolution of plants and microbes underpins sustainable agriculture. Microorganisms. 2021; 9(5), 1036.

63. Pyke GH. Plant-pollinator co-evolution: It’s time to reconnect with Optimal Foraging Theory and Evolutionary Stable Strategies. Perspect. Plant. Ecol. Evol. Syst. 2016; 19: 70–76.

64. Occhipinti A. Plant coevolution: evidences and new challenges. J. Plant Interact. 2013; 8(3): 188–196.

65. Wicaksono WA, Cernava T, Berg C, Berg G. Bog ecosystems as a playground for plant-microbe coevolution: bryophytes and vascular plants harbour functionally adaptive bacteria. Microbiome. 2021; 9(1), 170.

66. Aplet GH. & Laven RD. Relative performance of four Hawaiian shrubby plants (Asteraceae) under greenhouse conditions with implications for rarity. Biol. Conserv. 1993; 65(1): 15–21.

67. Walck JL, Baskin JM, Baskin CC. Relative competitive abilities and growth characteristics of a narrowly endemic and geographically widespread solidago species (Asteraceae). Am. J. Bot. 1999; 86(6): 820–828.

68. Lloyd KM, Lee WG, Wilson JB. Competitive Abilities of Rare and Common Plants: Comparisons Using *Acaena* (Rosaceae) and *Chionochloa* (Poaceae) from New Zealand. Conserv. Biol. 2002; 16(4): 975–985.

69. Brown KS. & Gilbert B. Population- and community-level rarity have opposing effects on pollinator visitation and seed set. J. Ecol. 2020; 108(5): 1835–1844.

70. Broz AK, Broeckling CD, De-la-Peña C, Lewis MR, Greene E, Callaway RM, et al. Plant neighbor identity influences plant biochemistry and physiology related to defense. BMC Plant Biol. 2010; 10, 115.

71. Hahn PG. & Orrock JL. Neighbor palatability generates associational effects by altering herbivore foraging behavior. Ecology. 2016; 97(8): 2103–2111.

72. Luo W, Lan R, Chen D, Zhang B, Xi N, Li Y, et al. Limiting similarity shapes the functional and phylogenetic structure of root neighborhoods in a subtropical forest. New Phytol. 2020; 229(2): 1078–1090.

73. Markham J. Rare species occupy uncommon niches. Sci. Rep. 2014; 4, 6012.

74. Alexander JM, Atwater DZ, Colautti RI, Hargreves AL. Effects of species interactions on the potential for evolution at species’ range limits. Philos. Trans. R. Soc. B: Biol. Sci. 2022; 377(1848), 20210020.

75. Mendham DS, Sankaran KV, O’Connell AM, Grove TS. *Eucalyptus globulus* harvest residue management effects on soil carbon and microbial biomass at 1 and 5 years after plantation establishment. Soil Biol. Biochem. 2002; 34(12): 1903–1912.

76. Forrester DI, Bauhus J, Cowie AL. Carbon allocation in a mixed-species plantation of *Eucalyptus globulus* and *Acacia mearnsii*. For. Ecol. Manag. 2006; 233(2-3): 275–284.

77. Forrester DI, Theiveyanathan S, Collopy JJ, Marcar NE. Enhanced water use efficiency in a mixed *Eucalyptus globulus* and *Acacia mearnsii* plantation. For. Ecol. Manag. 2010; 259(9): 1761–1770.

